# Global prevalence and phylogeny of hepatitis B virus (HBV) drug and vaccine resistance mutations

**DOI:** 10.1101/2020.10.07.329391

**Authors:** Jolynne Mokaya, Tetyana I Vasylyeva, Eleanor Barnes, M. Azim Ansari, Oliver G Pybus, Philippa C Matthews

## Abstract

**Introduction:** Vaccination and anti-viral therapy with nucleos(t)ide analogues (NAs) are key approaches to reducing the morbidity, mortality and transmission of hepatitis B virus (HBV) infection. However, the efficacy of these interventions may be reduced by the emergence of drug resistance-associated mutations (RAMs) and/or vaccine escape mutations (VEMs). We have assimilated data on the global prevalence and distribution of HBV RAMs/VEMs from publicly available data and explored the evolution of these mutations.

**Methods:** We analysed sequences downloaded from the Hepatitis B Virus Database, and calculated prevalence of 41 RAMs and 38 VEMs catalogued from published studies. We generated maximum likelihood phylogenetic trees and used treeBreaker to investigate the distribution of selected mutations across tree branches. We performed phylogenetic molecular clock analyses using BEAST to estimate the age of mutations.

**Results:** RAM M204I/V had the highest prevalence, occurring in 3.8% (109/2838) of all HBV sequences in our dataset, and a significantly higher rate in genotype C sequence at 5.4% (60/1102, p=0.0007). VEMs had an overall prevalence of 1.3% (37/2837) and had the highest prevalence in genotype C and in Asia at 2.2% (24/1102; p=0.002) and 1.6% (34/2109; p=0.009) respectively. Phylogenetic analysis suggested that most RAM/VEMs arose independently, however RAMs including A194T, M204V and L180M formed clusters in genotype B. We show evidence that polymorphisms associated with drug and vaccine resistance may have been present in the mid 20^th^ century suggesting that they can arise independently of treatment/ vaccine exposure.

**Discussion:** HBV RAMs/VEMs have been found globally and across genotypes, with the highest prevalence observed in genotype C variants. Screening for the genotype and for resistant mutations may help to improve stratified patient treatment. As NAs and HBV vaccines are increasingly being deployed for HBV prevention and treatment, monitoring for resistance and advocating for better treatment regimens for HBV remains essential.

## INTRODUCTION

Anti-viral therapy with nucleos(t)ide analogue (NA) agents is a central approach to reducing morbidity, mortality and transmission of hepatitis B virus (HBV) infection. NAs are used to suppress viraemia, thus reducing inflammatory liver damage (1). However, the efficacy of widespread deployment of NAs, both for individual patients and at a public health level, may be affected by the emergence of drug resistance (2,3). Resistance-associated mutations (RAMs) can arise as a result of the low fidelity of the HBV reverse transcriptase (RT) enzyme which lacks transcriptional proofreading activity, especially in the setting of high viral replication rates (estimated at up to ∼10^12^ virions/day (2)). Lamivudine (3TC) and entecavir (ETV) were licensed in 1986 and 2005, respectively, but their ongoing role has been limited by the occurrence of anti-viral drug resistance (4–6). Tenofovir (TFV), most commonly prescribed as tenofovir disoproxil fumarate (TDF), was licensed in 2008 and is now the favoured choice as it has a higher genetic barrier to resistance, as well as being cheap, well-tolerated, and safe, including in pregnancy (4). However, there are now emerging data that show the potential for selection of TDF drug resistance mutations (7), albeit with limited insights into their prevalence or clinical impact (8). Importantly, as well as being selected in individuals on therapy, RAMs have been reported among treatment-naïve individuals (1,9). Whether these mutations occur without exposure to antivirals, or are exclusively as result of prior drug exposure, is uncertain.

Reports of resistance to the HBV vaccine raise concerns about the extent to which vaccine-mediated immunity will remain robust. The vaccine, licensed for use in 1981, is administered to infants as part of WHO expanded programme for immunization (10). HBV vaccination induces the production of neutralising antibodies that mainly target the second hydrophilic loop (amino acids (aa) 139 to 147 or 149) of the major antigenic determinant (aa 99 to 169) of the HBV surface protein (HBsAg) (11,12). Strong immune pressure can lead to the selection of mutations within HBsAg, resulting in variants resistant to HBV vaccine and/or HBV immunoglobulin (HBIg) (2). G145A/R is the best described mutation associated with resistance to HBV vaccine/HBIg (11–13). Several other mutations across the entire antigenic determinant have been reported, which also have associations with vaccine resistance (11,14–16).

Genetic differences among the ten HBV genotypes (A-J) and numerous sub-genotypes may influence the likelihood of acquisition of drug or vaccine resistance (17). Genotypes have different geographical distributions, for example genotypes A, D and E are predominant in Africa, and B and C in Asia (18,19). In genotypes in which the wild type amino acid at a specific position is part of a sequence motif associated with drug or vaccine resistance, the barrier to resistance is likely to be inherently lower. This phenomenon has been described in hepatitis C virus (HCV) infection, explaining why some sub-genotypes are intrinsically resistant to the most widely used direct acting antiviral drugs (20–22). In addition, genotype-specific differences in mutation rates and host population dynamics have an influence on virus evolutionary rates, which directly affects the probability of appearance of RAMs/VEMs. For HBV, the rate of molecular evolution is estimated to be between 7.9 × 10^−5^ and 3.2 × 10^−4^ substitutions per site per year (23,24).

A number of studies have reported the frequencies of RAMs in HBV from different populations (1,25–27); however, the global prevalence, geographic distribution, time of origins, and their association with different HBV genotypes remain unknown. We therefore set out to assimilate data on the global prevalence and distribution of HBV RAMs from public sequence databases, and to explore the genetic relatedness of viruses bearing these mutations.

## METHODS

### HBV sequences curation process

We analysed sequences downloaded from a publicly available database (Hepatitis B Virus Database - https://hbvdb.ibcp.fr/HBVdb/ (28)), accessed on 20^th^ November 2018. We downloaded a total of 6,219 full length genome sequences (**Suppl Figure 1**). Using MEGA7 software (29), we generated neighbour joining phylogenetic trees to validate the HBV genotype assignment, discarding sequences that had been incorrectly classified. We then generated pairwise distances for aligned sequences within each genotype using the dist.alignment function of the R package seqinr (30), and excluded sequences with >99.5% similarity in order to remove closely related isolates for instance duplicates and/or isolates derived from the same individual. For the remaining sequences, we obtained sample collection date and sampling country from GenBank. A total of 2938 sequences had geographical data and 2167 had both sample collection date and geographical data. **Suppl Figure 1** shows the data curation process.

### Drug resistance associated mutations

We worked from a list of pre-existing drug RAMs identified from published studies (1,2,8,9,25,31) (**Suppl Table 1**), and stratified them according to the NA to which they cause resistance, as described below:

a. We classified RAMs associated with 3TC into three categories: (i) primary RAMs, which are well known to cause resistance to 3TC in isolation; (ii) compensatory RAMs, which by themselves do not confer resistance but when combined with primary RAMs enhance resistance and viral functional capacity (2); and (iii) putative RAMs for which there is limited clinical/phenotypic evidence for 3TC resistance.
b. Two or more amino acid substitutions are required across the HBV RT protein to confer resistance to ETV which could occur as a combination of M204I/V with one or more of the following substitutions L80I/V, I163V, I169T, V173L, L180M, A181S/T/V, T184X, A186T,S202C/G/I/R, M250I/V and/or C256S/G.
c. We classified RAMs associated with TFV into three categories: those with both clinical and *in vitro* evidence; those with only phenotypic evidence; and those with only experimental evidence, as described in a systematic literature review (8).

### Vaccine escape mutations

All pre-existing VEMs included in this study were identified from published studies (1,14–16,32–39) (**Suppl Table 2**). G145A/R and K141E/I/R have the strongest evidence base of clinical and *in vitro* data to support HBV vaccine resistance (33), while other VEMs are considered putative, as they are supported by less robust data.

### Prevalence analyses

For the global prevalence analysis, we included HBV sequences with known country of origin from genotypes A - E; we excluded genotypes F, G & H from this analysis because of low sample size (<100), resulting in a total of 2,838 sequences. For all polymorphisms that have been reported in association with resistance listed in **Suppl Table 1 and 2**, we calculated the prevalence as total number of sequences with a specified mutation out of the total number of sequences in each genotype/continent.

We carried out prevalence analysis reporting confidence intervals and p-values for individual RAMs common to 3TC, ETV and TFV, for individual or combined RAMs associated with ETV and TFV resistance, and for individual VEMs. We calculated confidence intervals using an online software Epitools (http://epitools.ausvet.com.au). We used Chi square test to compare the prevalence of RAMs/VEMs between different genotypes and between different continents

We used GraphPad Prism v7.0 for data visualisation and statistical analyses.

### Distribution of selected RAMs and VEMs on maximum likelihood phylogenetic trees

We generated maximum likelihood (ML) phylogenetic trees for HBV genotype A (n=290), B (n=730), C (n=1102), D (n=565) and E (n=150) sequences for which geographic information was available. We used the general time reversible nucleotide substitution model with gamma-distributed among-site rate variation (GTR + G) in IQ-TREE (40). We chose this model as it incorporates different rates for every nucleotide change and different nucleotide frequencies, thus allowing for most flexibility allowing us to avoid a model-selection step (41). We rooted the trees using the mid.point function of the R package phangorn (42).

For this analysis, we considered a total of 12 RAMs (S106C/G, D134E, R153W/Q, V173L, L180M, A181T/V, A194T, A200V, M204I/V, L217R, L229V/W, I269L). These RAMs were selected because they are primary RAMs or have robust evidence in causing resistance to 3TC, ETV and/or TFV. We also considered eight VEMs (C139S, S/T140I, P142S, S/T143L/M, D144A/E/G/N, G145A/E/R, K141A/I/R and C147S) which are located within the epitope (aa139 – 147) which is a neutralising epitope for the HBV vaccine.

We used treeBreaker (43) to test if sequences with individual mutations were randomly distributed across the branches of the phylogenetic trees reconstructed for each genotype. The program calculates per-branch posterior probability of having a change in the distribution of a discrete character and gives a Bayes factor to show the strength of this evidence (43). A Bayes factor of >30 indicates strong evidence that sequences with RAMs/VEMs are not randomly distributed on a phylogenetic tree. We performed this analysis for each mutation separately.

### Phylogenetic dating

We performed phylogenetic dating to estimate the times of emergence of mutations of interest, focused on RAMs V173L, L180M and M204I/V as they are well known to cause (individually or synergistically) resistance to 3TC, ETV and TDF (8), and VEMs G145A/R and K141E/I/R as they have been best described to cause HBV vaccine resistance (11–13). In this analysis we included genotypes that had >50 sequences with associated sampling date information: genotype A (n=170), B (n=594), C (n=906), D (n=336) and E (n=88). We manually inspected sequences for misalignments in AliView program (43) and then excluded codon positions associated with resistance (we excluded all sites listed in **Suppl Tables 1 and 2**) to ensure that parallel evolution RAMs/VEMs does not affect the phylogeny (44). We used ML phylogenetic trees generated for each genotype as described above and then dated these phylogenetic trees using IQ-TREE v2.0.3 (45). We resampled the trees 100 times and chose lognormal relaxed molecular clock model because it has performed best in other studies of HBV evolution (46,47). We used TempEst to estimate the molecular clock signal in our datasets by regressing the root-to-tip genetic distance of each sequence in the tree and its sampling date (48). Based on application of TempEst, we estimated the correlation between the dates of the tips of the sequences and the divergence from the root to be 7.8 × 10^−2^, 3.9 × 10^−1^, 4.3 × 10^−2^, 2.3 × 10^−2^ and 2.1x 10^−1^ for genotypes A, B, C, D and E, respectively. Due to the lack of correlation, we used the substitution rate estimated before (24,49) and therefore we thus fixed the mean substitution rate to 5.0 × 10^−5^ (SD 4.12 × 10^−6^) subs/site/year for all genotypes in all subsequent analyses. We reported the time to most recent common ancestor (TMRCA) of two or more sequences that clustered together having the same mutation as this TMRCA likely corresponds to the lower bound of the age of the mutation.

We also performed molecular clock phylogenetic analyses using Bayesian Evolutionary Analysis Sampling Trees (BEAST). This method has been described in **Suppl Methods**.

## RESULTS

### i. Global prevalence of HBV drug RAMs

We assessed the prevalence of polymorphisms associated with drug resistance across 48 different sites within RT protein in a total of 2838 full length HBV sequences, **Suppl Table 1**. 90% (43/48) of the sites had polymorphisms associated with drug resistance, **Suppl Fig 2**. 60% of the sites had polymorphisms associated with drug resistance occurring at the prevalence of between 0-10% in both genotypes and continents. Genotypes A and C, as well as Europe had the highest number of sites (nine sites for genotype A and C, and 11 sites for Europe) with polymorphisms associated with drug resistance occurring at a prevalence of >20%, **Suppl Fig 2**.

#### RAMs common to 3TC, ETV and/or TDF

RAMs L80I/M/V, V173L, L180M, A181T/V, T184X are common to 3TC, ETV and/or TFV. M204I/V had the highest prevalence at 3.8% (109/2838) **(Figure 1A)**. Genotype C had the highest prevalence of all of these mutations, apart from L80I/M/V which is most common in Genotype D (although not statistically significant) **(Figure 1B)**. L180M and M204I/V were both present in all genotypes and continents analysed in this manuscript **(Figure 1B, C)**. However, there were no significant differences in prevalence of these RAMs across continents.

**Figure 1:**
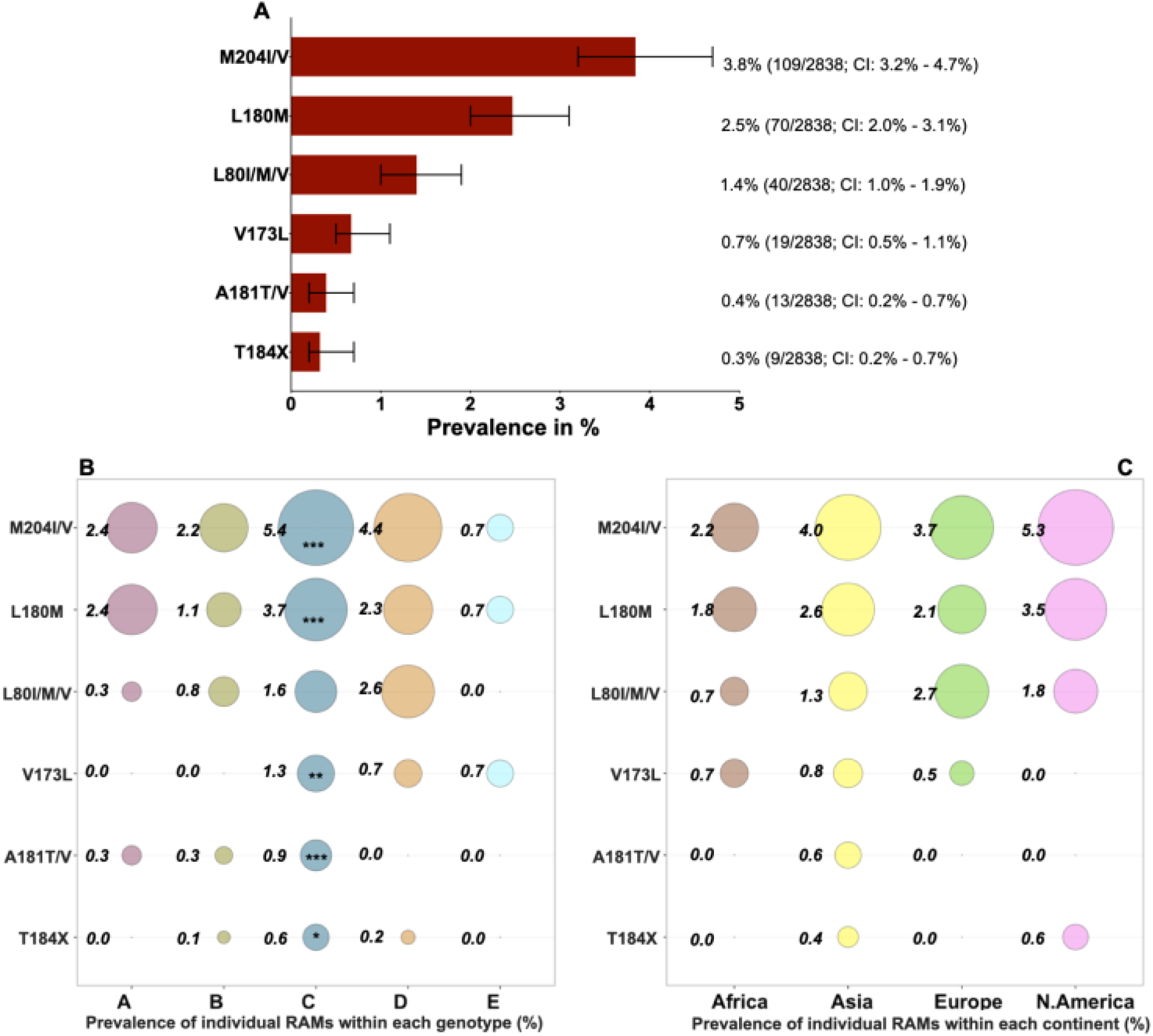
Global prevalence of hepatitis B virus (HBV) drug resistance associated mutations (RAMs) obtained from analysing 2838 HBV sequences with information on country of origin, downloaded from a public database (https://hbvdb.ibcp.fr/HBVdb/). **A**. Overall prevalence of RAMs common to 3TC, ETV and TFV. **B**. A bubble plot showing the overall prevalence of RAMs common to 3TC, ETV and TFV within each genotype (genotype A n=290; Genotype B n=730; Genotype C n=1102; Genotype D n=566; Genotype E n=150). **C**. A bubble plot showing the overall prevalence of RAMs common to 3TC, ETV and TFV within each continent (Africa n=277; Asia n=2109; Europe n=187; North America n=170). Numbers next to the circles are prevalence (%) of individual RAMs in each genotype/continent. The asterisks (***/**/*) within certain circles indicate RAMs that have a higher prevalence within the specified genotype/continent compared to the prevalence of that RAM in other genotypes/continents and is statistically significant. *** p value <0.001; **p value < 0.005; *p value <0.05. Bars show 95% confidence intervals. T184X represents T184A/C/F/G/I/L/M/S.

#### RAMs associated with ETV resistance

The overall prevalence of ETV resistance in this dataset, determined by the presence of RAMs M204I/V+L180M, was 2.4% (67/2838); other combinations of ETV drug resistant mutations were uncommon (all <0.6%); **Suppl Fig 3**. As previously, the most common resistance mutations were seen in genotype C at 3.5% (39/1102 vs 28/1736 in other genotypes; p=0.001).

#### RAMs associated with TFV resistance

The prevalence of individual mutations that have been associated with TFV resistance ranged between 0.2 – 19.5%. Compared to all other genotypes, genotype C had the highest prevalence of individual RAMs S106C/G, DH/N134E and I269L; and Asia had the highest prevalence of these individual RAMs S106C/G, DH/N134E and I269L compared to other continents, **Suppl Fig 4**.

Sequences with certain combinations of RAMs are likely to have the highest probability of clinically significant TFV resistance (8). We therefore sought evidence of these combinations of mutations in our sequence database (n=2838). In each case, we only identified between one and three sequences with each combination of RAMs giving an overall prevalence of between 0.04% - 0.1% (**Suppl Table 3**), suggesting these arise infrequently and are currently unlikely to be of significance at a population level. The majority of sequences carrying these drug resistance motifs were again in genotype C.

### ii. Global prevalence of VEMs

We assessed for the prevalence of polymorphisms associated with vaccine/ HBIg escape across 33 different sites within surface protein in a total of 2838 full length HBV sequences, **Suppl Table 2**. 78% (25/33) sites had polymorphisms associated with vaccine/HBIg escape, **Suppl Fig 5**. 52% (12/23) of the sites had polymorphisms associated with vaccine escape occurring at the prevalence of between 0 - 9% in both genotypes and continents. Genotype C and Asia had the highest number of sites with polymorphisms associated with vaccine escape occurring at the prevalence of >20%, compared to other genotypes and continents, **Suppl Fig 5**.

VEM K141E/I/R was not present in our dataset. G145A/R had an overall prevalence of 1.3% (37/2837) and had the highest prevalence in genotype C at 2.2% (24/1102; p=0.002), and in Asia at 1.6% (34/2109; p=0.009); this is the best recognised VEM **(Figure 2A)**. Other VEMs that had an overall prevalence of >1% were T118A/R/V, M133I/L/T, A128V, Q129H/N/R, G145A/R, P120S/T and S/T143L/M **(Figure 2A)**. T118A/R/V, A128V and S/T143L/M had the highest prevalence in genotype D and in Europe, being present >3% of the sequences; whereas VEMs M133I/L/T and Q129H/N/R had the highest prevalence in genotype B and M133I/L/T had the highest prevalence in Asia, also being present in >3% of the sequences **(Figure 2B and 2C)**.

**Figure 2:**
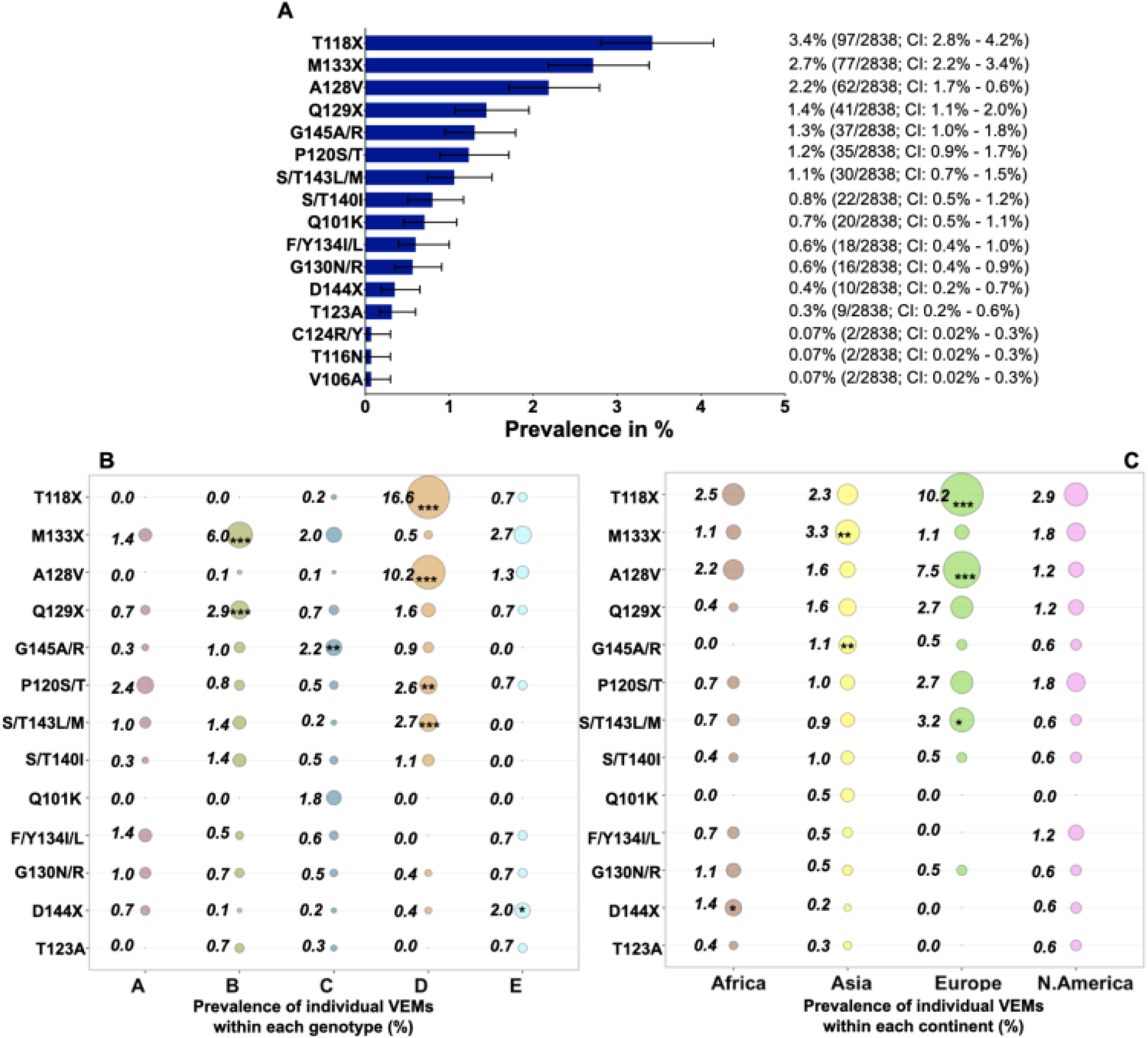
Global prevalence of hepatitis B virus (HBV) vaccine escape mutations (VEMs) obtained from analysing 2838 HBV sequences with information on country of origin, downloaded from a public database (https://hbvdb.ibcp.fr/HBVdb/). **A**. Overall prevalence of putative VEMs and/or VEMs with only clinical or *in vitro* evidence. **B**. A bubble plot showing the overall prevalence of putative VEMs and/or VEMs with only clinical or *in vitro* evidence within each genotype (genotype A n=290; Genotype B n=730; Genotype C; n=1102; Genotype D n=566 and Genotype E n=150), with a prevalence of >0.1%. **C**. A bubble plot showing the overall prevalence of putative VEMs and/or VEMs with only clinical or *in vitro* evidence within each continent (Africa n=277; Asia n=2109; Europe; n=187 and North America n=170), with a prevalence of >0.1%. Numbers next to the circles are prevalence (%) of individual RAMs in each genotype/continent. The asterisks (***/**/*) within certain circles indicate RAMs that have a higher prevalence within the specified genotype/continent compared to the prevalence of that RAM in other genotypes/continents and is statistically significant. *** p value <0.001; **p value <0.005; *p value< 0.05. T118X represents T118A/R/V; M133X represents M133I/L/T; Q129X represents Q129A/R; D144X represents D144A/E/G/N.

#### RAMs/VEMs as wildtype amino acid

Determining the clinical significance of individual RAMs/VEMs in HBV sequences is difficult because some of the mutations that have been described occur at consensus level in some genotypes. For example, 11 polymorphisms associated with drug resistance and nine polymorphisms associated with vaccine escape had a prevalence of >50% in ≥ 1 genotype (s), **Suppl Table 4** and **Suppl Table 5**. RAM H/Y9H is wildtype in genotypes A-E. This mutation is most likely to add to resistance as flexible positions in the protein, in which compensatory change is easily incorporated. RAMs H126Y and R/W153W, which contribute to TFV resistance when combined with ≥ 3 other RAMs (8), are wildtype in genotype A. This shows that resistance to different drugs or HBV vaccine might be more easily selected in certain populations or regions, based on the global distribution of HBV genotypes.

#### Distribution of selected RAMs and VEMs on maximum likelihood phylogenetic trees

We considered the distribution of 12 RAMs (S106C/G, D134E, R153W/Q, V173L, L180M, A181T/V, A194T, A200V, M204I/V, L217R, L229V/W, I269L) and eight VEMs (C139S, S/T140I, P142S, S/T143L/M, D144A/E/G/N, G145A/E/R, K141A/I/R and C147S) across the branches of ML phylogenetic trees. Most of these RAMs and all VEMs were randomly distributed across the branches of phylogenetic trees reconstructed from genotypes A-E sequences, suggesting parallel evolution.

However, there were several RAMs that clustered within genotype B, C and D sequences (**Figures 3)**. In genotype B, all sequences containing the A194T variant clustered together (Bayes factor, BF, support >100; n=4 sequences). Sequences with this RAM were all from Indonesia, reported by a study exploring HBV genetic diversity (50). Some sequences containing both M204V and L180M formed a cluster in genotype B (BF = 54.99, n=4 sequences) and some with M204I formed a cluster in genotype D (BF >100, n=3 sequences). In genotype C, there were clusters of RAMs S106C (BF >100, n=5 sequences), R153Q (BF >100, n=3 sequences), and I129L (BF >100, n=34 sequences). Clustering of sequences with RAMs might suggest an emerging sublineage of treatment resistant virus.

**Figure 3:**
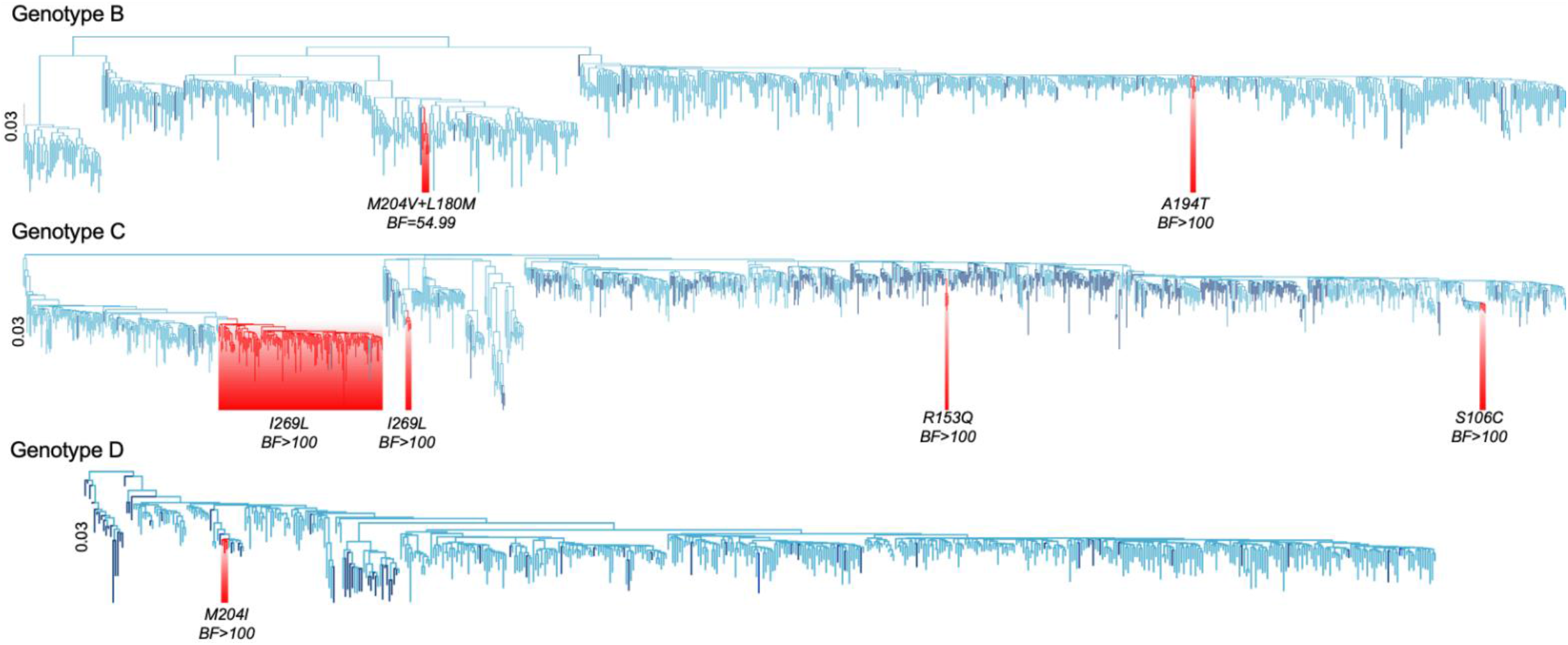
HBV RAMs/VEMs distribution on rooted maximum likelihood phylogenetic trees for genotype B, C and D. Branches in dark blue represent sequences with one or more RAMs/VEMs. Branches in light blue have no specified RAMs/VEMs. Branches highlighted in red indicate clustered sequences with a RAM with Bayes factor of >30, suggesting strong evidence of clustering. ML trees for genotype A and E were not displayed because they had no sequences with specified RAM/VEM which formed clusters.

#### Evolution of sequences with RAMs/VEMs

Most sequences with RAMs/VEMs in our analysis were published after the approval of NAs/HBV vaccine, as a result of widespread improvements and availability of sequencing that have arisen in parallel with roll out of drugs and vaccine. However, four sequences (KF214668, KF214671, KF214673 and KF214676) with RAM I269L and one sequence (KF214659) with VEM S/T143M were sampled from Asia in 1963, and one sequence (HQ700441) with RAM L180M was sampled from Oceania in 1984, demonstrating that mutations can arise without exposure to treatment or vaccination.

We performed ML molecular clock analysis for full datasets of sequences of genotypes A – E. However, only genotype C had clusters that had at least two isolates with the same resistant mutations with a single common ancestor as shown in **Table 1**. The estimated time of emergence of branches with RAMs M204V+L180M was around year 1945 (95% HPD 1897 - 1971); and branches with VEM G145R was estimated to emerge around year 1930 (95% HPD 1866 - 1958). Importantly, in both cases the higher bound of the 95% HPD interval of the TMRCA of these clusters, which likely correspond to the lower bound of the estimate of the age of these mutations, precedes the introduction of NAs and the HBV vaccine. The results we obtained from ML molecular clock analysis and BEAST analysis were consistent.

**Table 1:** Estimated time of the most common recent ancestor (TMRCAs) (and 95% HPD) of branches with specific RAMs/VEMs on molecular clock trees. HPD: Highest Posterior Density. Only genotype C is displayed because it had at least two isolates with the same resistant mutations with a single common ancestor.

| Genotype | RAMs/VEMs | Cluster of isolates with specified RAMs/VEMs | Estimated TMRCAs (95%HPD) |
| --- | --- | --- | --- |
| C | M204V+L180M | FJ032355 | 1945 (1897, 1971) |
|  |  | FJ386620 |  |
|  | G145R | KU964229 | 1930 (1866,1958) |
|  |  | KU964230 |  |

## DISCUSSION

### Novel findings and comparison with previous literature

We describe the global prevalence of drug and vaccine resistance in HBV across genotypes and geographical regions and explore the evolution of these mutations using phylogenetic analysis, in order to provide a high-resolution picture of the origins and distribution of drug resistance. From this analysis, HBV drug and vaccine resistance are not common, with the highest frequency of individual or combined mutations that are well known to cause resistance, being ∼4% and the majority being <1%. These mutations are distributed across various continents and genotypes, with the most frequent RAMs/VEMs identified in genotype C, concordant with previous studies from China (51,52). We show that these mutations are not only driven by exposure to drug or vaccine but are likely to have been present in some sequences for hundreds of years. More studies representing all genotypes are needed, alongside careful correlation with clinical evidence of drug resistance.

M204I/V is one of the best recognised drug resistance motifs in HBV and had the highest overall prevalence of 3.8% within our whole sequence database. A previous meta-analysis estimated the prevalence of M204I/V as 4.9% among >12,000 treatment-naïve individuals (25), and another review reported a prevalence of M204I/V of 5.9% among 8, 435 treatment-naïve individuals (9). These reviews reported prevalence as the proportion of individuals with mutations within the total number of treatment naïve individuals, without accounting for closely related sequences (thus may include multiple sequences from a single individual). In contrast, we used full length HBV sequences and excluded identical sequences, which might explain the lower prevalence we report.

We reported the prevalence of ETV resistance as 2.4%, which is slightly higher than the prevalence of 1.7% reported from a large survey carried out in China among 1223 treatment experienced patients and also a prevalence of 1.2% reported from a longitudinal study that followed 108 HBV infected treatment naïve individuals for five years (26,53). Unlike these two studies, we took a lenient approach by reporting the overall prevalence of ETV resistance considering sequences with RAMs M204I/V+L180M, with or without an additional compensatory mutation. These two are always present in ETV resistant variants, and are the main ETV RAMs reported in published HBV treatment guidelines (10,54).

We estimated the overall prevalence of TFV resistance to be between 0.04 - 0.2%. There have been few studies that have reported on TFV resistance (8) and more robust data are still needed to define HBV resistance in order to guide better estimation of the prevalence of relevant RAMs.

The global prevalence of the VEM G145A/R in our data was 1.3%, which is comparable to that be 0.3 - 1% previously reported across genotypes A-F (55). A study carried out in Italy reported a higher prevalence of 3.1%, in a cohort dominated by genotype D infection (56). Regional differences might explain the difference in prevalence. However, the majority of individuals with this mutation from the Italian study were immunocompromised and it was not clear if they had been vaccinated prior to becoming infected.

### Relationship between genotype and drug or vaccine resistance

The prevalence of RAMs/VEMs across different regions is influenced by the predominant genotype, but may also relate to different patterns of drug or vaccine exposure in the population. For example, T118A/R/V, A128V and D144A/E/G/N variants are more common in Europe, which may relate to better vaccine coverage (57) that drives the selection of resistant variants. Some polymorphisms that have been described in association with resistance are wildtype in certain genotypes, which might indicate that these genotypes are more susceptible to the development of clinically significant drug resistance. For example, TFV resistance might be selected more easily in genotype A given that RAMs H126Y and R/W153W are wildtype in this genotype (58).

### Phylogenetic analysis of selected RAMs and VEMs

We provide evidence that RAMs can arise without exposure to treatment/vaccine, showing that certain RAMs emerged prior to the NAs and vaccine era. Using phylogenetic dating, we estimate that RAMs M204V and L180M, and VEM G145R were already present around the mid 20^th^ century. Although these estimates have wide confidence intervals, their upper bounds precede the time of introduction of NAs and the HBV vaccine. A previous study estimated the origin date of HBV genotype D in Iran as 1894 (95% HPD 1701 – 1957)(47), and the root age of genotype A polymerase sequences is estimated as the year 955 (95% HPD 381 – 1482); (46). A study that analysed 167 full length genotype E sequences, estimated the TMRCA to be 174 years (95% HPD 36 – 441); (59). Similar to our analysis, these studies used an uncorrelated relaxed lognormal clock which is reported to the best fitting clock (46,47,59). However, given the differences in the substitution models, and with some studies using sequences for just a single gene, direct comparison of the estimated TMRCA generated by these studies and our analysis is challenging.

### Selection vs transmission of drug resistance

Most RAMs were randomly distributed across the branches of HBV phylogenetic trees, which suggests that these polymorphisms are being selected independently in individual hosts (parallel evolution (60)) rather than becoming fixed and disseminated from a founder strain. The high viral replication and mutation rate of HBV can result in amino acid substitutions at sites of resistance, leading to the stochastic emergence of drug RAMs even in individuals who have not been exposed to treatment (61–63). Individuals can also be infected with HBV strains containing drug RAMs which could significantly comprise virological response to therapy, as has been shown in HIV (64).

### Caveats and limitations

The major constraint in this work is the relative lack of HBV sequence data; given the huge global burden of infection there is a striking lack of high-quality sequence data available in the public domain. As our sequences were obtained from GenBank, metadata on individual characteristics and treatment exposure were not available. Our analyses may not be representative, given the biased nature of sequence data that are available, disproportionately representing certain populations and regions, and samples containing high viral load (65). Drug resistant sequences may be over-represented, given that virus suppressed by drug therapy is not accessible for sequencing and individuals with break-through viraemia on treatment are more likely to have samples submitted for sequence analysis.

Phylogenetic dating in HBV is challenging. Its overlapping reading frame raises controversies around its evolution rates. HBV sequences lack temporal signal thus making it challenging to reliably date HBV evolution using molecular clock methods. In addition, estimation of TMRCA uses sample collection dates obtained from GenBank, which may not be accurate.

While Asia and Africa are known to have the highest prevalence of chronic HBV infection worldwide at 6.2 and 6.1% respectively (66), 74% (2109/2838) of sequences included in this analysis were from Asia and only 10% (277/2838) published from Africa. This low representation of sequences highlights HBV as a neglected disease, with very few individuals diagnosed and linked to care (67). In addition, the influence of the widespread use of antiretroviral drugs containing 3TC and TFV, on suppression and/or emergence of drug resistance is not yet understood.

## Conclusions

Despite the availability of effective prevention and treatment strategies for HBV infection, emergence of RAMs and VEMs may pose a challenge to the achievement of the United Nations sustainable development goals for elimination by 2030. Going forward, enlarged sequencing datasets, collected together with treatment histories and clinical data, will be essential to develop an understanding of the distribution, nature and significance of drug resistance at an individual and population level.

## AUTHOR CONTRIBUTIONS

Conceived the study: JM, PCM. Assimilated data: JM, MAA. Analysed the data: JM, TIV, MAA. Wrote the manuscript: JM, TIV, PCM. Revised the manuscript: JM, TIV, EB, MAA, OP, PCM. All authors have read and approved the manuscript.

## Supplementary Figures

**Suppl Fig 1:**
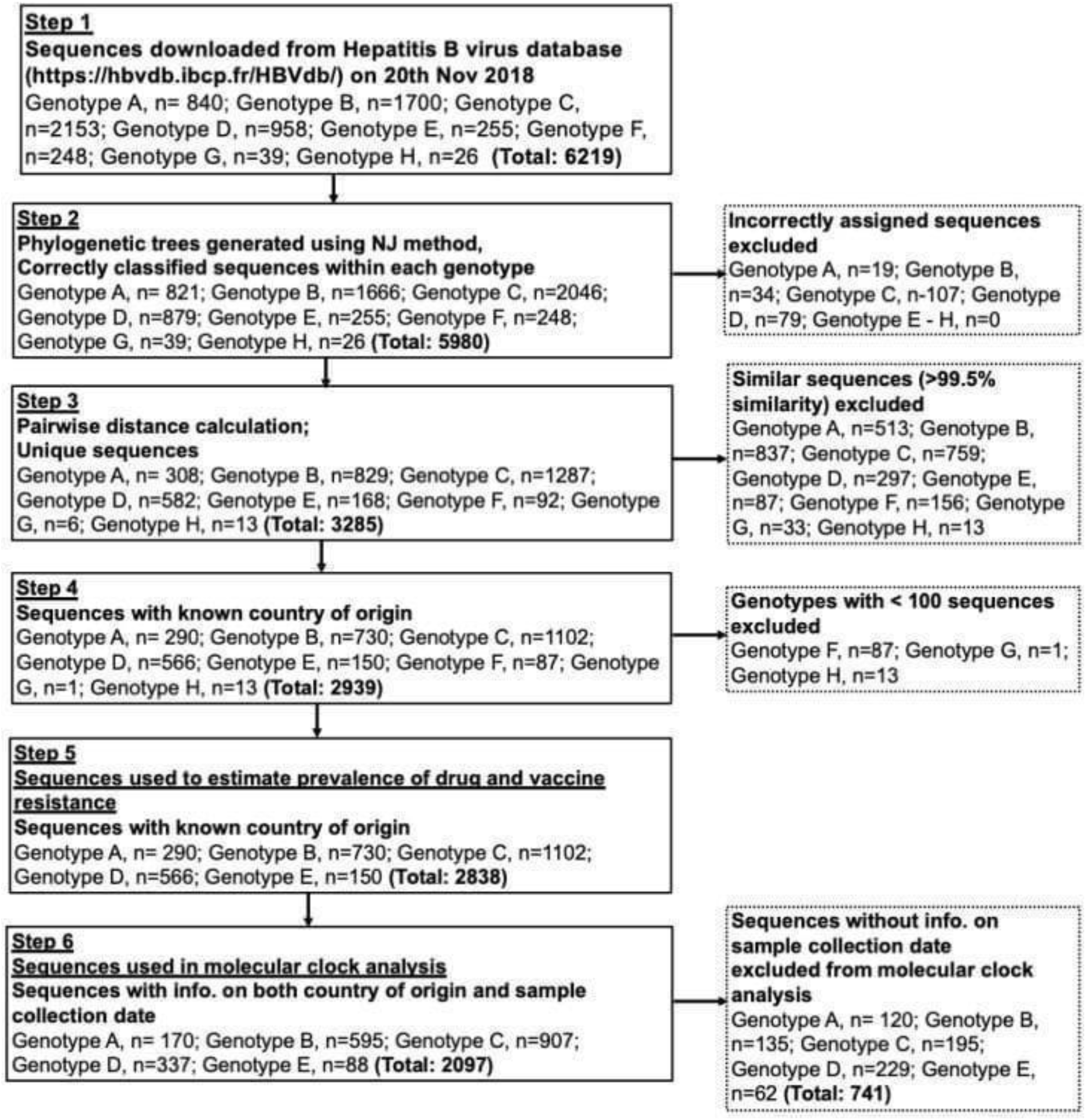
Flow diagram showing data curation process of sequences downloaded from a public database (https://hbvdb.ibcp.fr/HBVdb/) included in the analysis of the global prevalence and evolution of hepatitis B virus (HBV) drug resistance associated mutation (RAMs) and vaccine escape mutations (VEMs).

**Suppl Fig 2:**
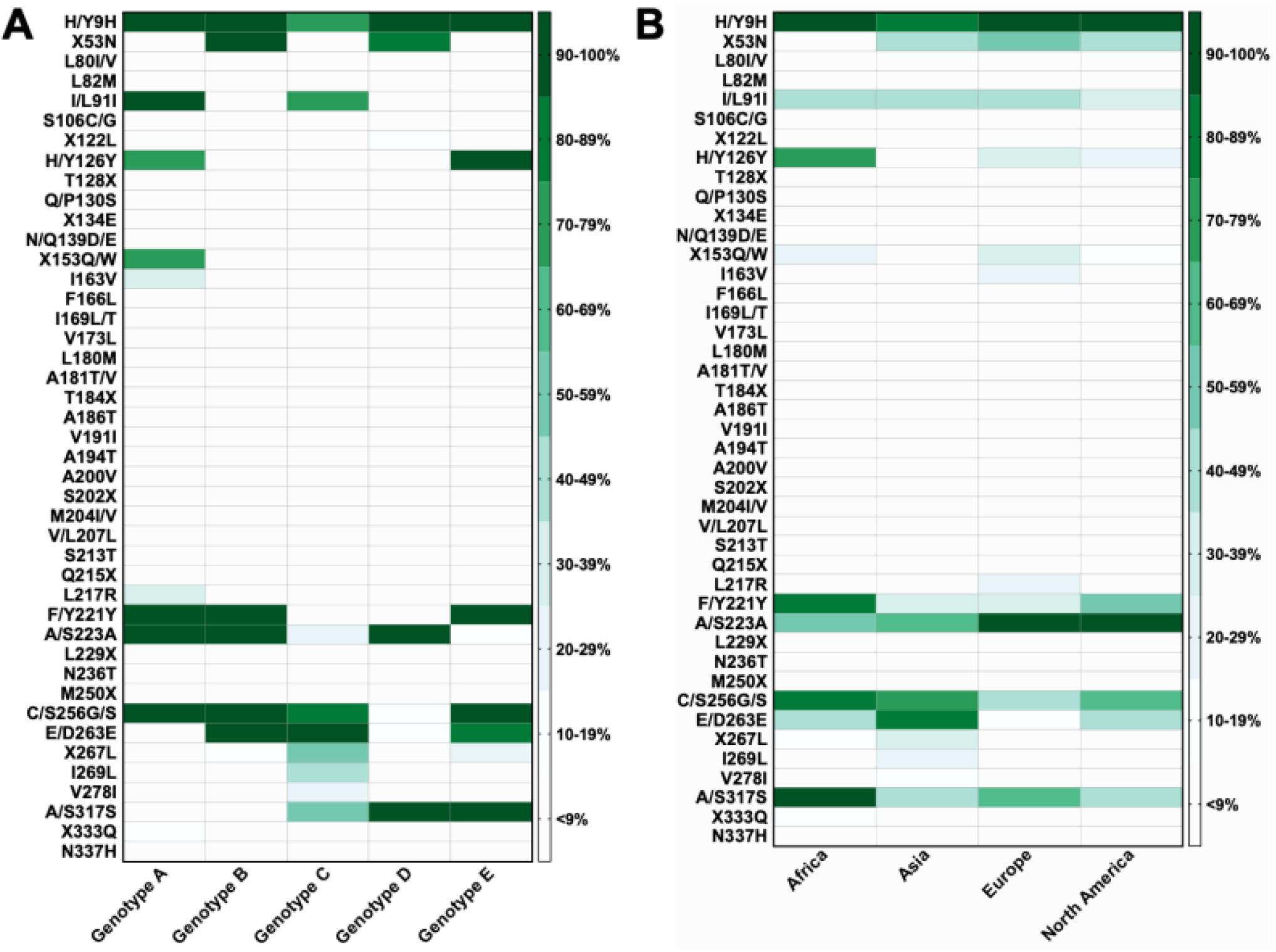
Global prevalence of hepatitis B virus (HBV) drug resistance associated mutations (RAMs) obtained from analysing 2838 HBV sequences with information on country of origin, downloaded from a public database (https://hbvdb.ibcp.fr/HBVdb/). **A**. Prevalence of polymorphisms across genotypes; **B**. Prevalence of polymorphisms across continents. X53N represents V/N/S/T53N; X122L represents I/F/H/L/N/Y122L; T128X represents T128A/I/N; X134E represents D/H/N134E; X153Q/W represents Q/R/W153Q/W; T184X represents T184A/C/F/G/I/L/M/S; S202X represents S202C/G/I; Q215X represents Q215E/H/P/S; L229X represents L229G/F/V/W; M250X represents M250I/L/V; X267L represents H/L/M/Q267L.

**Suppl Fig 3:**
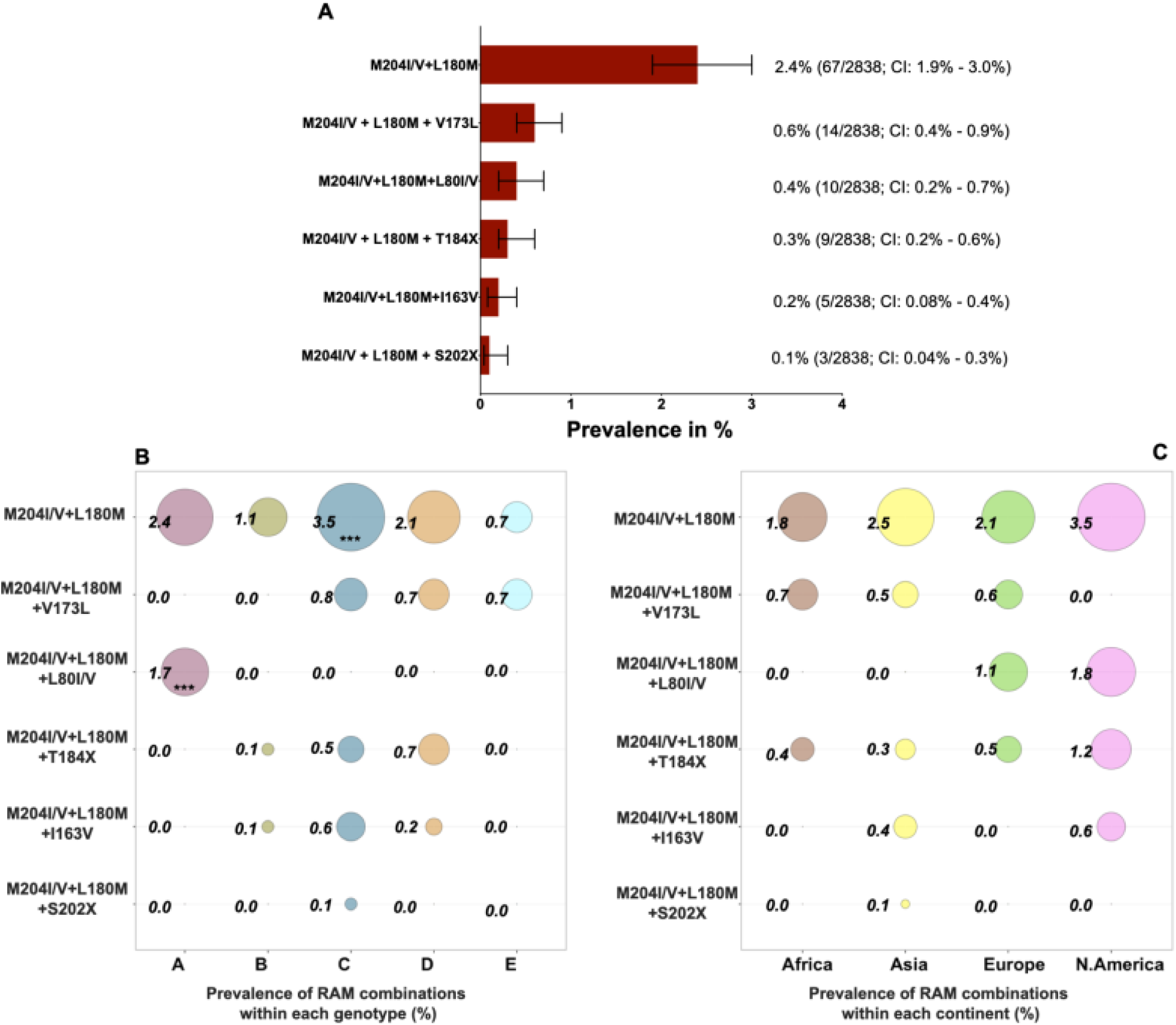
Global prevalence of hepatitis B virus (HBV) entecavir (ETV) resistance associated mutations (RAMs) obtained from analysing 2838 HBV sequences with information on country of origin, downloaded from a public database (https://hbvdb.ibcp.fr/HBVdb/). **A**. Overall prevalence of ETV RAMs. **B**. A bubble plot showing the prevalence of ETV RAMs within each genotype (genotype A n=290; Genotype B n=730; Genotype C; n=1102; Genotype D n=566 and Genotype E n=150). **C**. A bubble plot showing the prevalence of ETV RAMs within each continent (Africa n=277; Asia n=2109; Europe; n=187 and North America n=170). Numbers next to the circles are prevalence (%) of individual RAMs in each genotype/continent. The asterisks (***/**/*) within certain circles indicate RAMs that have a higher prevalence within the specified genotype/continent compared to the prevalence of that RAM in other genotypes/continents and is statistically significant. *** p value <0.001; **p value < 0.005; *p value <0.05. Bars show 95% confidence intervals. T184X represents T184A/C/F/G/I/L/M/S and S202X represents S202C/G/I/R

**Suppl Fig 4:**
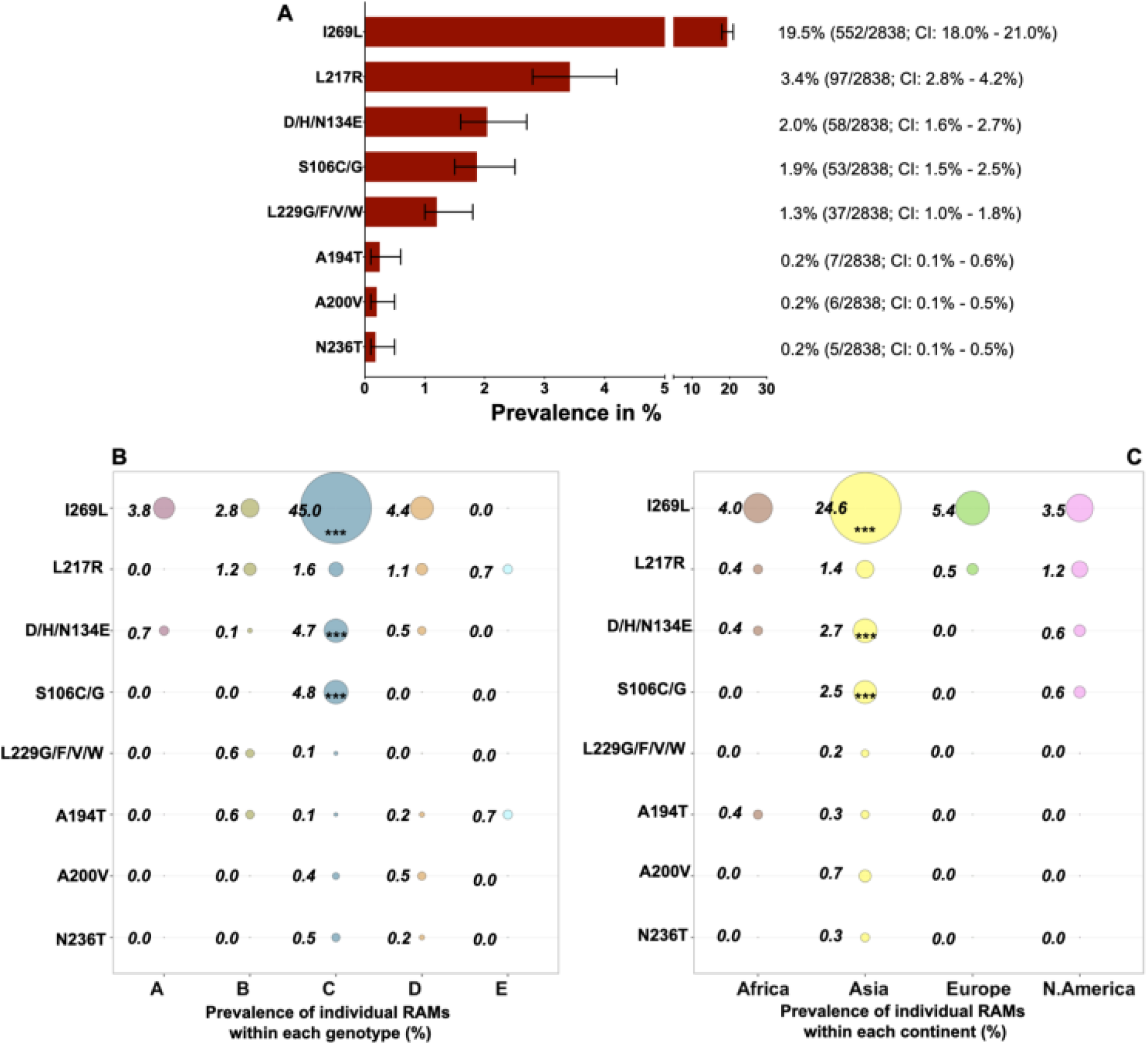
Global prevalence of hepatitis B virus (HBV) tenofovir (TFV) resistance associated mutations (RAMs) obtained from analysing 2838 HBV sequences with information on country of origin, downloaded from a public database (https://hbvdb.ibcp.fr/HBVdb/). **A**. Overall prevalence of TFV RAMs. **B**. A bubble plot showing the overall prevalence of TFV RAMs within each genotype (genotype A n=290; Genotype B n=730; Genotype C; n=1102; Genotype D n=566 and Genotype E n=150). **C**. A bubble plot showing the overall prevalence of TFV RAMs within each continent (Africa n=277; Asia n=2109; Europe; n=187 and North America n=170). Numbers next to the circles are prevalence (%) of individual RAMs in each genotype/continent. The asterisks (***/**/*) within certain circles indicate RAMs that have a higher prevalence within the specified genotype/continent compared to the prevalence of that RAM in other genotypes/continents and is statistically significant. *** p value <0.001; **p value < 0.005; *p value <0.05. Bars show 95% confidence intervals.

**Suppl Fig 5:**
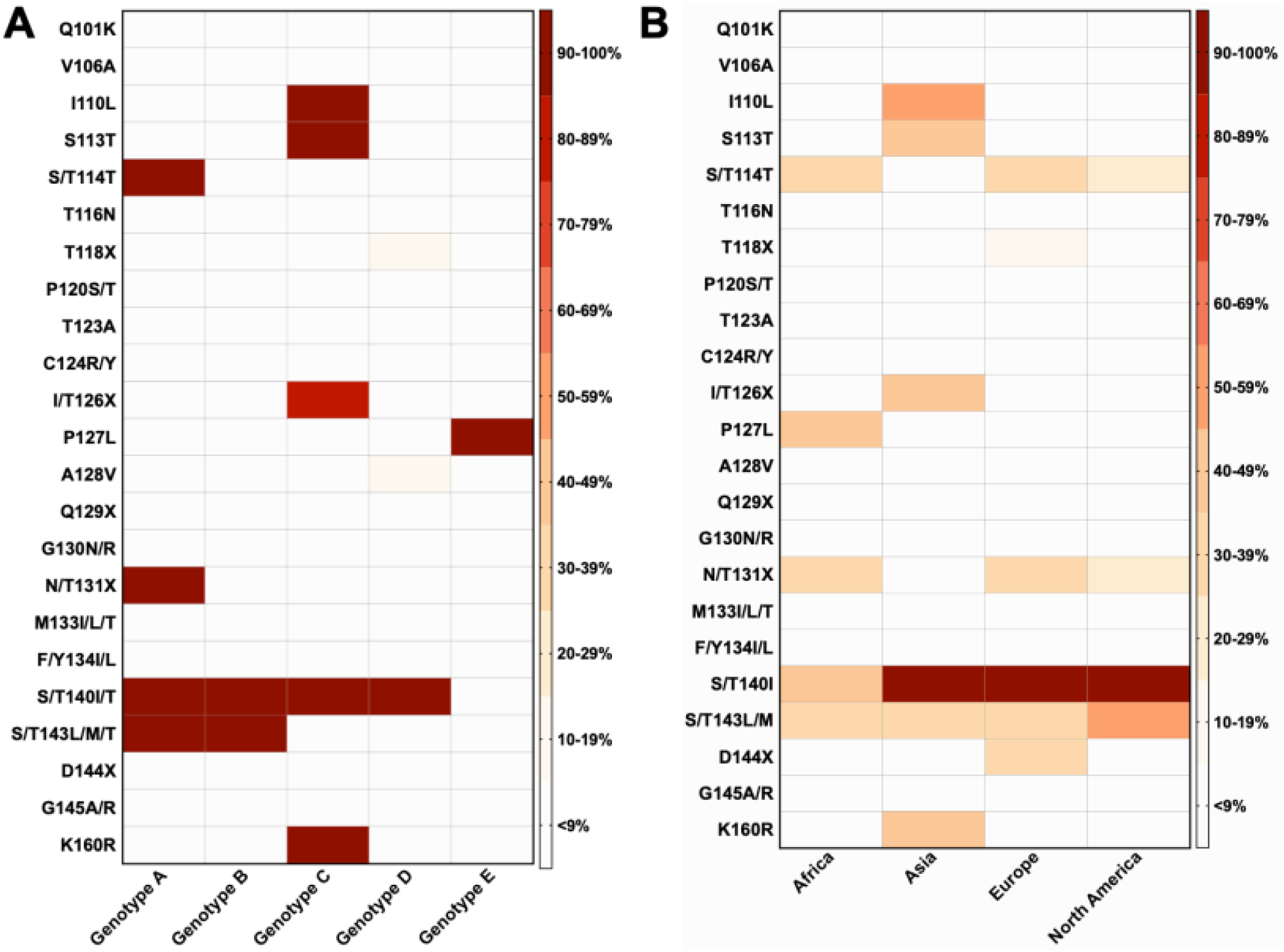
Global prevalence of hepatitis B virus (HBV) vaccine escape mutations (VEMs) across genotypes, obtained from analysing 2838 HBV sequences with information on country of origin, downloaded from a public database (https://hbvdb.ibcp.fr/HBVdb/) **A**. Showing prevalence of polymorphisms across genotypes; **B**. Showing prevalence of polymorphisms across continents.

## Supplementary Tables

**Suppl Table 1:**
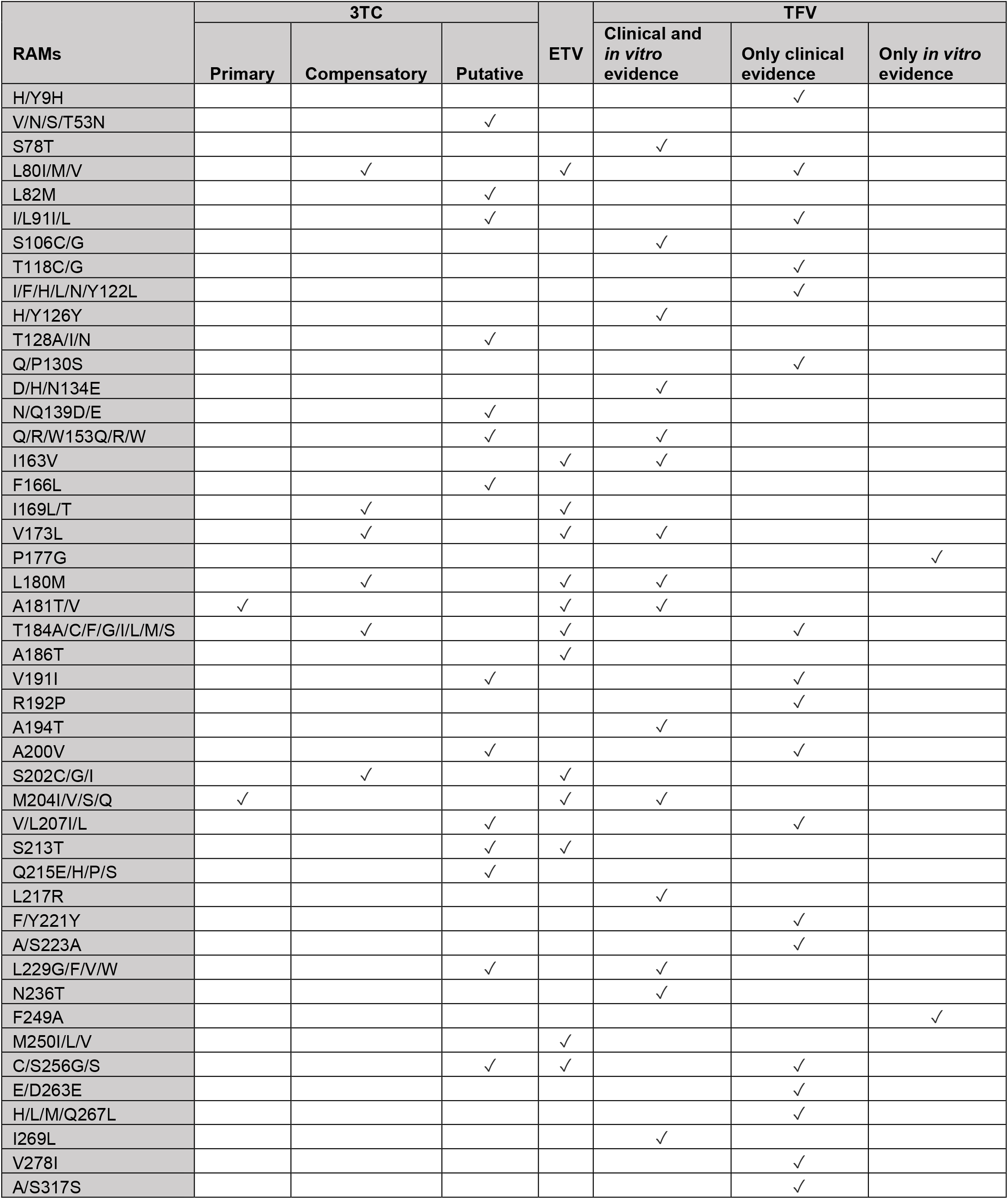
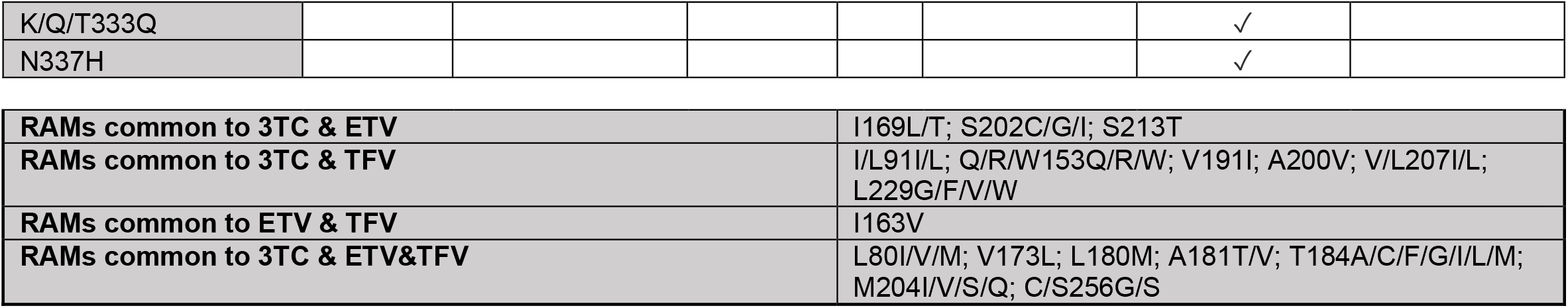
Hepatitis B virus drug resistance associated mutations (RAMs). Data obtained from published systematic reviews (1,2,8,9,25,31). Amino acid positions listed in HBV reverse transcriptase protein. 3TC: Lamivudine; ETV: Entecavir; TFV: Tenofovir

| RAMs | 3TC |  |  | ETV | TFV |  |  |
| --- | --- | --- | --- | --- | --- | --- | --- |
|  | Primary | Compensatory | Putative |  | Clinical and *in vitro* evidence | Only clinical evidence | Only *in vitro* evidence |
| H/Y9H |  |  |  |  |  | ✓ |  |
| V/N/S/T53N |  |  | ✓ |  |  |  |  |
| S78T |  |  |  |  | ✓ |  |  |
| L80I/M/V |  | ✓ |  | ✓ |  | ✓ |  |
| L82M |  |  | ✓ |  |  |  |  |
| I/L91I/L |  |  | ✓ |  |  | ✓ |  |
| S106C/G |  |  |  |  | ✓ |  |  |
| T118C/G |  |  |  |  |  | ✓ |  |
| I/F/H/L/N/Y122L |  |  |  |  |  | ✓ |  |
| H/Y126Y |  |  |  |  | ✓ |  |  |
| T128A/I/N |  |  | ✓ |  |  |  |  |
| Q/P130S |  |  |  |  |  | ✓ |  |
| D/H/N134E |  |  |  |  | ✓ |  |  |
| N/Q139D/E |  |  | ✓ |  |  |  |  |
| Q/R/W153Q/R/W |  |  | ✓ |  | ✓ |  |  |
| I163V |  |  |  | ✓ | ✓ |  |  |
| F166L |  |  | ✓ |  |  |  |  |
| I169L/T |  | ✓ |  | ✓ |  |  |  |
| V173L |  | ✓ |  | ✓ | ✓ |  |  |
| P177G |  |  |  |  |  |  | ✓ |
| L180M |  | ✓ |  | ✓ | ✓ |  |  |
| A181T/V | ✓ |  |  | ✓ | ✓ |  |  |
| T184A/C/F/G/I/L/M/S |  | ✓ |  | ✓ |  | ✓ |  |
| A186T |  |  |  | ✓ |  |  |  |
| V191I |  |  | ✓ |  |  | ✓ |  |
| R192P |  |  |  |  |  | ✓ |  |
| A194T |  |  |  |  | ✓ |  |  |
| A200V |  |  | ✓ |  |  | ✓ |  |
| S202C/G/I |  | ✓ |  | ✓ |  |  |  |
| M204I/V/S/Q | ✓ |  |  | ✓ | ✓ |  |  |
| V/L207I/L |  |  | ✓ |  |  | ✓ |  |
| S213T |  |  | ✓ | ✓ |  |  |  |
| Q215E/H/P/S |  |  | ✓ |  |  |  |  |
| L217R |  |  |  |  | ✓ |  |  |
| F/Y221Y |  |  |  |  |  | ✓ |  |
| A/S223A |  |  |  |  |  | ✓ |  |
| L229G/F/V/W |  |  | ✓ |  | ✓ |  |  |
| N236T |  |  |  |  | ✓ |  |  |
| F249A |  |  |  |  |  |  | ✓ |
| M250I/L/V |  |  |  | ✓ |  |  |  |
| C/S256G/S |  |  | ✓ | ✓ |  | ✓ |  |
| E/D263E |  |  |  |  |  | ✓ |  |
| H/L/M/Q267L |  |  |  |  |  | ✓ |  |
| I269L |  |  |  |  | ✓ |  |  |
| V278I |  |  |  |  |  | ✓ |  |
| A/S317S |  |  |  |  |  | ✓ |  |
| K/Q/T333Q |  |  |  |  |  | ✓ |  |
| N337H |  |  |  |  |  | ✓ |  |

|  |  |
| --- | --- |
| **RAMs common to 3TC & ETV** | I169L/T; S202C/G/I; S213T |
| **RAMs common to 3TC & TFV** | I/L91I/L; Q/R/W153Q/R/W; V191I; A200V; V/L207I/L;<br>L229G/F/V/W |
| **RAMs common to ETV & TFV** | I163V |
| **RAMs common to 3TC & ETV&TFV** | L80I/V/M; V173L; L180M; A181T/V; T184A/C/F/G/I/L/M;<br>M204I/V/S/Q; C/S256G/S |

**Suppl Table 2:** Hepatitis B virus vaccine escape mutations (VEMs). Data obtained from published studies (1,14–16,32–39). Amino acid positions listed in HBV surface protein. VEM: Vaccine escape mutation. HBsAg: Hepatitis B surface antigen.

| VEMs | HBIG | VEMs | Immune escape |
| --- | --- | --- | --- |
| Q101K |  |  | ✓ |
| V106A |  |  | ✓ |
| I110L |  |  | ✓ |
| S113T |  |  | ✓ |
| S/T114F/R/T | ✓ |  | ✓ |
| T116A/N |  | ✓ |  |
| T118A/R/V | ✓ |  |  |
| P120A/E/N/Q/S/T | ✓ | ✓ | ✓ |
| R/K122S |  |  |  |
| T123A/N | ✓ |  | ✓ |
| C124R/Y | ✓ |  |  |
| I/T126A/H/I/N/R/S | ✓ | ✓ | ✓ |
| P/T127L/P |  | ✓ | ✓ |
| A128V |  |  | ✓ |
| Q129H/N/R/P | ✓ | ✓ | ✓ |
| G130N/R | ✓ |  | ✓ |
| N/T131I/N/S | ✓ | ✓ | ✓ |
| M133I/L/T | ✓ | ✓ | ✓ |
| F/Y134I/L | ✓ | ✓ | ✓ |
| C137R/Y | ✓ |  |  |
| C138Y |  |  | ✓ |
| C139S | ✓ |  |  |
| S/T140I | ✓ |  | ✓ |
| K141E/I/R | ✓ | ✓ |  |
| P142S | ✓ | ✓ | ✓ |
| S/T143M/L |  | ✓ |  |
| D144A/E/G/H/N | ✓ | ✓ | ✓ |
| G145A/K/R | ✓ | ✓ | ✓ |
| N146S | ✓ |  | ✓ |
| C147S | ✓ |  |  |
| K160R |  |  | ✓ |

**Suppl Table 3:** Description of sequences with individual or combination of RAMs that are highly likely to cause resistance to TFV obtained from analysing 2838 HBV sequences with information on country of origin, downloaded from a public database (https://hbvdb.ibcp.fr/HBVdb/). These RAMs combination include ≥ 1 RAMs from the ‘short list’ in combination with ≥ 3 other RAMs from the ‘long list’ as described (8).

| Sequence ID | Individual and combination of mutations that are highly likely to cause resistance to TFV | No. of sequences with RAM (%) | Continent and number of sequences | Genotype and number of sequences |
| --- | --- | --- | --- | --- |
| GQ358144-7;<br>GQ377536;<br>KT366495;<br>GQ161771 | A194T* | 7 (0.2) | Asia n=6;<br>Africa n=1 | B, n=4;<br>C, n=1;<br>D, n=1,<br>E, n=1, |
| FJ386681 | A181T/V*+N236T | 1 (0.04) | Asia n=1 | B n=1 |
| KJ803809 | S106C*+D134E*+Q267L+I269L*+K333Q | 1 (0.04) | Asia n=1 | C n=1 |
| FJ386579;<br>FJ787470;<br>FJ787471 | S106C*+V173L*+L180M*+M204I/V*+Q267L | 3 (0.1) | Asia n=3 | C n=3 |
| EU939588 | S106C*+L180M*+A200V+M204I*+Q267L | 1 (0.04) | Asia n=1 | C n=1 |
| FJ386623 | S106C*+L180M*+ M204I/V*+Q267L | 1 (0.04) | Asia n=1 | C n=1 |
| FJ386620;<br>JX026877 | S106C*+L180M*+ M204I/V*+I269L | 2 (0.07) | Asia n=2 | C n=2 |
| AB182589 | S106C*+D134E*+Q267L+I269L*+K333Q+N3337H | 1 (0.04) | Asia n=1 | C n=1 |
| JQ040132 | D134E*+L180M*+M204I*+I267L+N337H | 1 (0.04) | Asia n=1 | C n=1 |
| AY641561 | D134E*+I267L*+K333Q+N337H | 1 (0.04) | Asia n=1 | C n=1 |
| EU560439 | D134E*+Q267L+I269L*+K333Q | 1 (0.04) | Asia n=1 | C n=1 |
| JN827423 | R153Q*+V173L*+L180M*+M204V*+I269L*+V278I | 1 (0.04) | Asia n=1 | C n=1 |
| JQ707346 | R153W*+L180M*+M204V*+V207L+L217R* | 1 (0.04) | North America<br>n=1 | A n=1 |
| FJ899789 | R153Q*+Q267L+I269L*+V173L*+N337H | 1 (0.04) | Asia n=1 | C n=1 |
| MF772345 | R153W*+L217R*+V278I+K333Q | 1 (0.04) | Africa n=1 | A n=1 |
| JN257203 | R153Q*+F122L+V278I+K333Q | 1 (0.04) | Africa n=1 | D n=1 |
| KX357637 | V173L*+L180M*+M204V*+V207L | 1 (0.04) | Asia n=1 | D n=1 |
| FJ032355 | V137L+L180M+M204V+Q267L+I269L* | 1 (0.04) | Asia n=1 | C n=1 |
| JN827418;<br>JN827421;<br>MF925409 | V137L*+L180M*+M204V*+I269L*+V278I | 3 (0.1) | Asia n=3 | C n=3 |
| AB697490 | V137L*+L180M*+M204V*+N337H | 1 (0.04) | Asia n=1 | C n=1 |
| FJ787453 | V137L*+ M204V*+ Q267L+I269L* | 1 (0.04) | Asia n=1 | C n=1 |
| FJ386604 | L180M*+A181V*+M204V*+I269L* | 1 (0.04) | Asia n=1 | C n=1 |
| JF828921 | L180M*+T184L+M204V*+L229V*+Q267L+K333Q+N337H | 1 (0.04) | Asia n=1 | C n=1 |
| JF828923;<br>JF828937 | L180M*+T184A/L+M204V*+ Q267L+K333Q+N337H | 2 (0.07) | Asia n=2 | C n=2 |
| EU939564; | L180M*+A200V+M204I*+Q267L | 1 (0.04) | Asia n=1 | C n=1 |
| FJ386653;<br>FJ787455;<br>FJ787456 | L180M*+M204I/V*+L229V*+Q267L | 3 (0.1) | Asia n=3 | C n=2 |
| JN827422;<br>JN827424 | L180M*+M204I/V*+I269L*+V278I | 2 (0.07) | Asia n=2 | C n=2 |
| DQ246215 | L180M*+M204I*+V278I+N337H | 1 (0.04) | Asia n=1 | C n=1 |

\* RAMs provided in 'short list' described in (8); these RAMs are supported by the highest quality evidence (i.e. isolated from treatment compliant individuals in whom viraemia was not suppressed by TFV and these RAMs were also tested in *in vitro* assays to measure the effect of TFV on viral replication in cell lines)

**Suppl Table 4:**
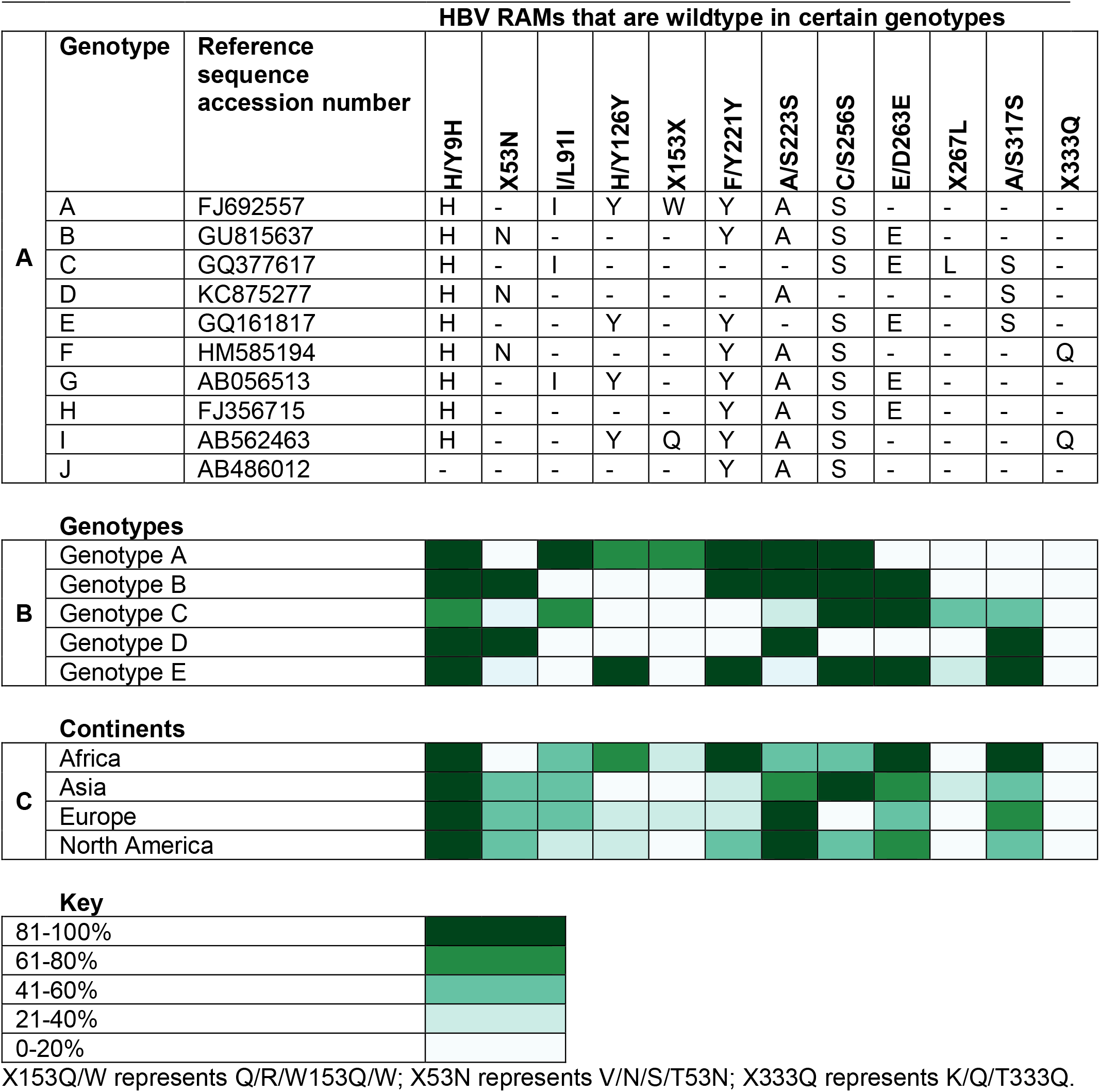
Global prevalence of hepatitis B virus (HBV) drug resistance associated mutations (RAMs) that are wildtype amino acid. Prevalence rates were obtained from analysing 2838 HBV sequences with information on country of origin, downloaded from a public database (https://hbvdb.ibcp.fr/HBVdb/). **A**. Identification of RAMs as wildtype in certain genotypes using HBV reference sequences for genotypes A-J. **B**. Prevalence of RAMs that are wildtype amino acid across genotypes. **C**. Prevalence of RAMs that are wildtype amino acid across continents. HBV reference sequences were obtained from a published manuscript (68).

**Suppl Table 5:**
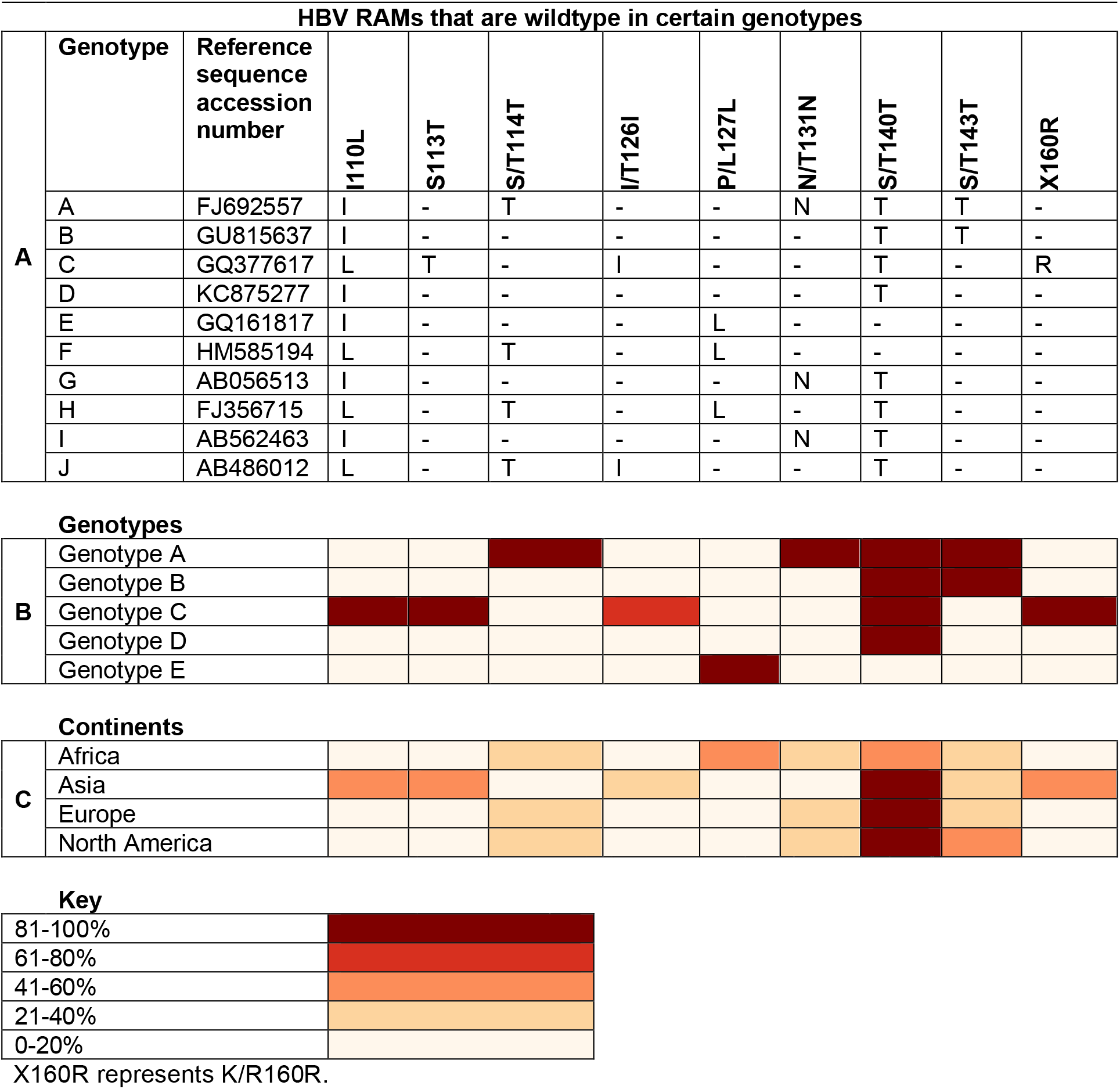
Global prevalence of hepatitis B virus (HBV) vaccine escape mutations (VEMs) that are wildtype amino acid. Prevalence rates were obtained from analysing 2838 HBV sequences with information on country of origin, downloaded from a public database (https://hbvdb.ibcp.fr/HBVdb/). **A**. Identification of VEMs as wildtype in certain genotypes using HBV reference sequences for genotypes A-J. **B**. Prevalence of VEMs that are wildtype amino acid across genotypes. **C**. Prevalence of VEMs that are wildtype amino acid across continents.

## Suppl Methods: Phylogenetic dating using Bayesian Evolutionary Analysis Sampling Trees (BEAST)

We performed molecular clock phylogenetic analyses to estimate the times of emergence of mutations of interest, focussed on RAMs V173L, L180M and M204I/V as they are well known to cause (individually or synergistically) resistance to 3TC, ETV and TDF (8), and VEMs G145A/R and K141E/I/R as they have been best described to cause HBV vaccine resistance (11–13). In this analysis we included genotypes that had >50 sequences with associated sampling date information: genotype A (n=170), B (n=594), C (n=906), D (n=336) and E (n=88). We manually inspected sequences for misalignments in AliView program (43) and then excluded codon positions associated with resistance (we excluded all sites listed in **Suppl Tables 1 and 2**) to ensure that parallel evolution RAMs/VEMs does not affect the phylogeny (44). We first identified sequences containing these mutations on the molecular clock tree and then only focused on reporting the time to most recent common ancestor (TMRCA) of two or more sequences that clustered together having the same mutation.

We performed Bayesian Markov chain Monte Carlo (MCMC) analyses using BEAST v.1.10 (69). We used a GTR+G nucleotide substitution model, a coalescent Bayesian Skygrid model with 50 points (70) and the uncorrelated lognormal relaxed molecular clock model. These models were selected because they have performed best in other studies estimating HBV evolution (46,47). TempEst allows quantification of temporal signal by estimating regressing the root-to-tip genetic distance of each sequence in the tree and its sampling date (48). Based on application of TempEst, we estimated the correlation between the dates of the tips of the sequences and the divergence from the root to be 7.8 × 10^−2^, 3.9 × 10^−1^, 4.3 × 10^−2^, 2.3 × 10^−2^ and 2.1x 10^−1^ for genotypes A, B, C, D and E, respectively, and thus we elected not to estimate the molecular clock rate as there was insufficient signal in our data. We thus fixed the mean substitution rate to 5.0 × 10^−5^ (SD 4.12 × 10^−6^) and a mean standard deviation of 2.0 × 10^−5^ (SD 4.96 × 10^−7^) subs/site/year, for all genotypes in all subsequent BEAST analyses, as this rate has been estimated before and applied in phylodynamic analyses of HBV (24,49).

To avoid convergence issues, we selected smaller subsamples from each of our alignments, depending on their size, to ensure that alignments used in BEAST analyses were <200 sequences. Thus, Genotype B (total n=564) was split into 3 subsamples, genotype C (total n=906) into 6 subsamples and genotype D (total n=336) into 2 subsamples. We used stratified random sampling, ensuring equal representation of sequences with mutations in each subsample. We ran one BEAST analysis for Genotypes A and E since their full alignments contained <200 sequences.

Two MCMC chains of 100 × 10^6^ generations (10% burn-in) were run with sampling every 10,000^th^ generation for genotypes A and E, and for each subsample of genotypes B, C and D separately. The MCMC chains for analyses of the same genotype were combined using LogCombiner v1.10.4 (69). We inspected convergence of the MCMC analyses using Tracer v.1.7.1 (71) to ensure effective sample size (ESS) >200 for all model parameters. We inferred maximum clade credibility trees using TreeAnnotator v.1.10.0 (69) and visualised them using FigTree v.1.4.4 (69).

